# Long noncoding RNA NRAV can regulate interferon-stimulated gene expression in melanoma

**DOI:** 10.64898/2026.08.25.746911

**Authors:** Kadir Ziya Durmus, Emre Kilic, Cagatay Sahin, Sude Ece Aral, Huseyin Atakan Ekiz

## Abstract

The long non-coding RNA Negative Regulator of Antiviral Response (NRAV) is known to suppress antiviral immunity by regulating interferon response, but its functional role in tumor immunology remains poorly understood. We examined NRAV’s relevance in melanoma and found that high NRAV expression was associated with poor survival, reduced inflammatory pathway activation, and resistance to immune checkpoint blockade. Bulk and single-cell transcriptomic profiling indicates that NRAV expression is selectively enriched in malignant cells suggesting a potential cancer cell-intrinsic function. To examine whether NRAV can regulate inflammatory responses in melanoma cells, we manipulated the levels of NRAV in the BRAF-mutant A375 melanoma model and characterized the expression of key interferon-stimulated genes (ISGs) following type-I and type-II interferon stimulation. Our findings reveal that the stable NRAV overexpression blunts the induction of key ISGs, whereas NRAV knockdown reciprocally amplifies their transcription. Subcellular fractionation revealed that NRAV is predominantly localized to the nuclear compartment of melanoma cells and the overexpression of NRAV altered regulatory histone marks on the target ISG promoters including MX1 and IFITM3. Collectively, these findings establish NRAV as a tumor-intrinsic epigenetic regulator of interferon signaling, highlighting its potential contribution to melanoma immune evasion.

## Introduction

Cutaneous melanoma is an aggressive skin malignancy that has served as a primary model for understanding cancer immunology and the clinical development of immune checkpoint inhibitors (ICIs) (1,2). While these ICIs have yielded durable clinical benefit in some patients, primary and acquired resistance remain a significant challenge to treatment failure in a large proportion of patients (3). This resistance is frequently driven by tumor-intrinsic genetic or epigenetic adaptations that allow malignant cells to evade host defenses. Consequently, identifying tumor-intrinsic pathways that govern resistance and dictate therapy responsiveness represents a critical clinical priority (3).

Within the tumor microenvironment (TME), signaling pathways activated by both Type-I (IFNα/β) and Type-II (IFNγ) interferons serve as crucial drivers of tumor clearance and therapeutic efficacy, acting through Janus kinase-signal transducer and activator of transcription (JAK-STAT) cascade to induce a conserved program of transcriptionally active interferon-stimulated genes (ISGs) (4). Several studies have reported that higher IFNγ pathway activity within the TME correlates with improved prognosis and enhanced immunotherapy response (4,5). Nevertheless, the role of interferons in tumor immunity is highly complex; while acute exposure is immunostimulatory, chronic or persistent signaling within the tumor can paradoxically coordinate adaptive resistance networks (6,7). Furthermore, tumor cell-intrinsic interferon signaling has been shown to induce regulatory feedback networks that alter chromatin states and immunomodulatory gene expression, underscoring the necessity of how malignant cell lineages dynamically tune their response to microenvironmental cytokines (8).

Beyond classic protein-coding transcriptional networks, long non-coding RNAs (lncRNAs) have emerged as key determinants of cellular differentiation, tissue homeostasis, and disease progression (9). LncRNA expression is highly cell-type and tissue-specific, and these molecules are frequently dysregulated in human cancers, acting as highly precise biomarkers or functional oncogenic drivers (10,11). LncRNAs execute their versatile regulatory roles through diverse mechanisms, such as forming molecular scaffolds to stabilize protein-protein interactions, acting as competing endogenous RNAs to sponge target transcripts, and directly interacting with chromatin-remodeling complexes and histone modifiers to alter the epigenetic landscape (9).

Among such regulatory lncRNAs is the Negative Regulator of Antiviral Response (NRAV) originally characterized in 2014 as a potent repressor of viral immunity in lung epithelial cancer models (12). Mechanistically, NRAV dampens inflammatory gene expression in the context of Type I interferon response and drives the accumulation of repressive H3K27me3 marks at the promoters of key ISGs, such as MxA and IFITM3. This study also reported that transgenic mice expressing human NRAV succumb to viral infection due to suboptimal antiviral immunity, suggesting NRAV regulates conserved immunological pathways in mice and humans. Beyond this antiviral setting, only a handful of studies have investigated the function of NRAV to date, demonstrating its capacity to interact with specific microRNAs and proteins to drive oncogenic signaling, metabolic reprogramming, and cell survival in other solid tumors (13–16). Together, these studies establish NRAV as a dynamic modulator of complex pro-tumorigenic epigenetic, post-transcriptional, and metabolic networks. Despite its emerging role as an immune and oncogenic regulator, NRAV’s function in melanoma remains unexplored. Given that infectious and tumor microenvironments share conserved interferon signaling pathways to coordinate immune responses, these findings provide a strong rationale to investigate NRAV in antitumor immunity.

In this study, we hypothesize that human melanoma cells can leverage epigenetic suppressor NRAV to selectively blunt interferon-induced gene induction, thereby constructing an epigenetic shield against immune pressure. By combining multi-cohort clinical transcriptomic profiling of patient tumors with single-cell RNA-sequencing (scRNA-seq) datasets, we first establish that NRAV expression is highly tumor-selective within the melanoma microenvironment and serves as a robust predictor of poor clinical outcomes and checkpoint blockade failure. Next, we systematically validate this hypothesis through gain- and loss-of-function models in human BRAF-mutant A375 melanoma cells, showing that NRAV levels reciprocally govern the early transcriptional induction of classical ISGs—specifically MX1, IFIT2, and IFIT3—following exposure to both Type I and Type II interferons. Finally, using subcellular fractionation and chromatin immunoprecipitation (ChIP-qPCR), we reveal that nuclear-localized NRAV drives an asymmetric chromatin remodeling process at target promoters, promoting an enrichment of repressive H3K27me3 over active H3K4me3 marks to enforce a bivalent poised or repressed promoter architecture. Together, our results uncover a previously unrecognized epigenetic axis of interferon refractoriness in melanoma, highlighting NRAV as a potential target to restore microenvironmental cytokine responsiveness.

## Results

NRAV was previously shown to inhibit type-I interferon signaling in epithelial cells (12), but its relevance in melanoma progression and tumor immunity has not been investigated. Using The Cancer Genome Skin Cutaneous Melanoma (TCGA-SKCM) RNA-seq data, we first explored whether tumors expressing different levels of NRAV are associated with differential survival outcomes. When the samples are categorized at the median NRAV expression, NRAV-high cohort exhibited significantly poorer clinical outcomes, which was paralleled by a 47% risk increase in the Cox proportional hazards (CoxPH) modeling **(Fig.1a)**. An inflamed tumor microenvironment in SKCM was shown to correlate with better overall survival (17). We hypothesized that, if NRAV was involved in dampening melanoma antitumor immunity, its higher expression would correlate with reduced survival of patients with inflamed tumors selectively. To address this, we examined survival based on two factors: NRAV expression, and the IFNγ/IFNα pathway scores as calculated by gene set variation analysis (GSVA) (18,19). In these analyses, IFNγ-high/NRAV-low group demonstrated the most favorable survival outcome, and the NRAV expression level did not strongly affect the survival in the IFNγ-low cohort **(Fig.1b)**. When other key clinical covariates are included in the multivariate CoxPH modeling, NRAV continued to be significantly associated with poor outcome in melanoma. Notably, the presence of IFNγ score in the model partially reduced the statistical significance of and the risk provided by NRAV **(Fig.1c)**, supporting the immune-related functions of NRAV. We observed similar trends when NRAV was faceted by IFNα score in survival analysis, while NRAV’s survival associations were affected less in multivariable analyses **(Fig.1d-e)**. Although IFNγ and IFNα cytokines have evolved distinct signal transduction mechanisms and phenotypic functions, they can regulate a common set of target genes (20). We found that 36% of IFNγ and 75% of IFNα response pathway genes are mutually shared, which explains similar patterns in survival analysis **(Fig.1f)**. Among the common genes are MX1, IFIT2, IFIT3 and IFITM3 which were shown to be regulated by NRAV in the context of antiviral defense (12). Taken together, NRAV arises as a poor prognosticator in melanoma which is potentially associated with the regulation of tumor immunity.

**Figure 1.**
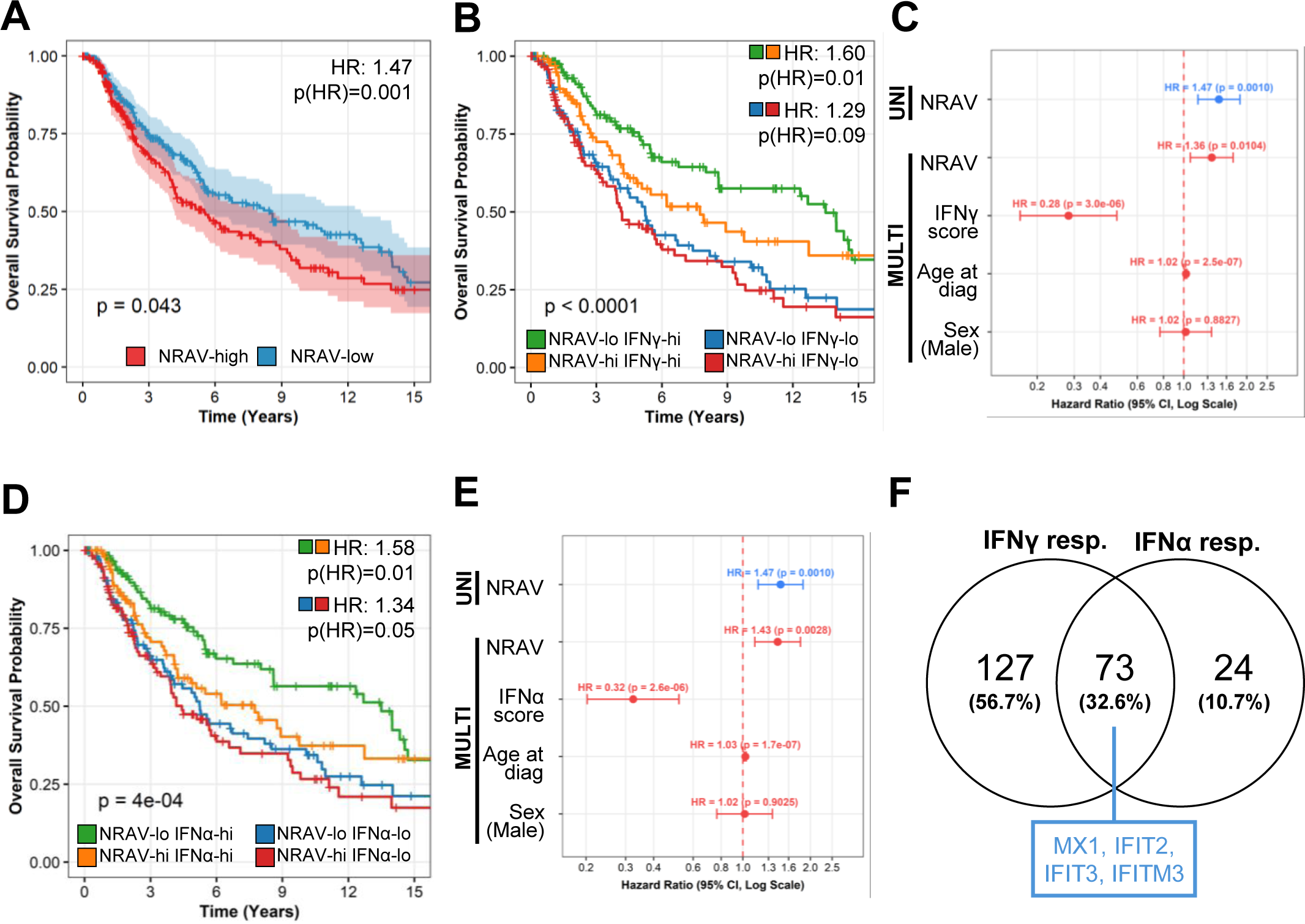
LncRNA NRAV is a poor prognosticator in human skin cutaneous melanoma (TCGA- SKCM). **A.** Higher NRAV expression within the TME is associated with shorter survival as evidenced by median-categorized Kaplan-Meier analysis and Cox proportional hazards (CoxPH) modeling. Log-rank and Wald test p-values are shown on the plot. **B.** Two-factor survival analysis incorporating NRAV expression and IFNγ score (GSVA) reveals that most favorable survival is attained when tumors have high IFNγ pathway activity and low NRAV expression. **C.** Forest plots from univariable (blue) and multivariable (red) CoxPH survival modeling demonstrate that NRAV is an independent predictor of poor clinical outcomes. **D.** Two- factor survival analysis involving NRAV and the IFNα pathway score demonstrates the most favorable clinical outcome in the NRAV-low/IFNα-high group. **E.** NRAV is a poor prognosticator in melanoma in multivariable CoxPH model including the IFNα score. **F.** Overlap analysis of Hallmark IFNα and IFNγ response gene sets indicates that a significant number of genes are shared.

After observing NRAV’s association with survival outcome in melanoma, we wanted to examine its relationship with the interferon-responsive gene expression and immunotherapy outcomes. Single-sample GSVA analysis of the TCGA-SKCM cohort revealed that IFNγ and IFNα responsive genes show highly concordant expression patterns, as expected due to partial overlap in the gene sets **(Fig.2a)**. Interestingly however, NRAV expression showed a stronger inverse relationship with the IFNγ response score compared to IFNα **(Fig.2b-c)**. To understand whether IFNγ/IFNα-responsive gene expression is differentially enriched in NRAV-dichotomized groups, we performed gene set enrichment analysis (GSEA) and observed that NRAV-low tumor samples express relatively higher levels of interferon pathway genes **(Fig.2d)**. IFNα has been historically used in melanoma treatment although it is now replaced by better immunotherapies (20), and the IFNγ responsive gene expression within the TME has been shown to predict immunotherapy outcomes (4,5). Thus, we next evaluated whether NRAV expression can also distinguish melanoma patients who benefits from nivolumab, an immune checkpoint inhibitor (ICI) targeting PD-1 (21). We noted that the NRAV expression tended to be lower in patients responding to treatment (complete or partial response, CR/PR) both prior to and during the treatment **(Fig.2e)**. Importantly, this trend was more readily observed during where both progressive and stable disease (PD/SD) exhibited significantly higher levels of NRAV expression. To evaluate the potential of NRAV in predicting unresponsiveness to nivolumab, receiver operating characteristic (ROC) curve analyses were performed comparing non-responders (PD/SD) with responders (CR/PR). Pre-treatment NRAV levels demonstrated moderate predictive capacity (AUC = 0.692), while on-treatment NRAV expression showed enhanced discriminatory performance (AUC = 0.744), indicating that dynamic changes in NRAV during therapy may provide superior predictive value for clinical response **(Fig.2f)**. The re- analyzed dataset from Riaz et al. comprises two distinct patient cohorts: an ipilimumab-refractory (prior progression on anti-CTLA-4 treatment) group and an ipilimumab-naïve group (21). We next evaluated whether NRAV expression associates with poor therapy response across these subsets and found that NRAV distinguished nivolumab non-responders with greater accuracy in the ipilimumab-refractory cohort **(Fig.2g)**. Together, these findings indicate that lncRNA NRAV can predict ICI responsiveness in melanoma.

**Figure 2.**
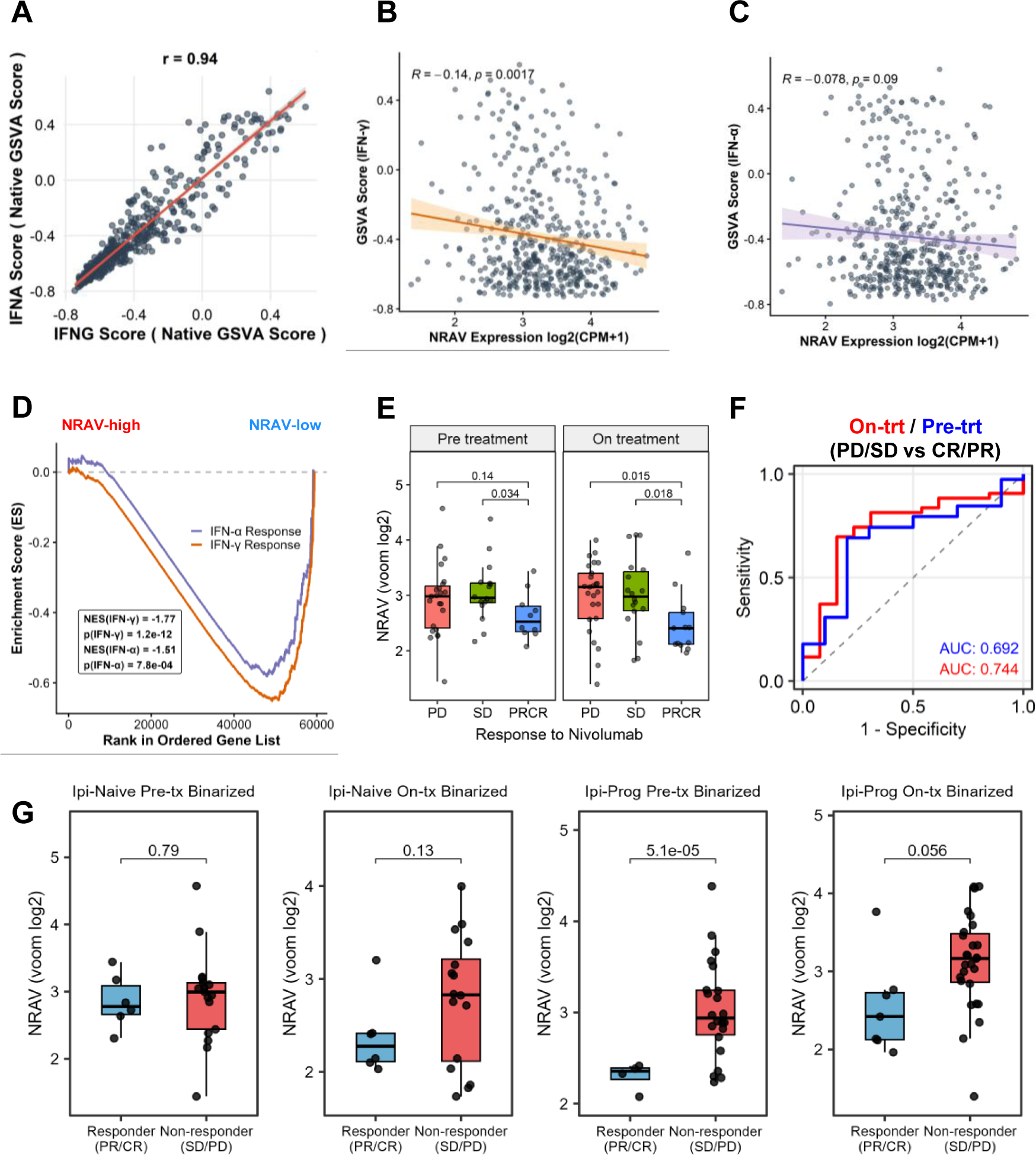
NRAV expression negatively correlates with interferon signaling and predicts poor response to immune checkpoint blockade in melanoma. **A.** Gene set variation analysis (GSVA) score calculated for IFNα and IFNγ response gene sets are plotted in the TCGA-SKCM RNA-seq samples. **B-C.** NRAV expression is negatively correlated with IFNγ (B) and IFNα (C) pathway scores, although this association was stronger for IFNγ pathway. **D.** Gene set enrichment analysis between NRAV-high and NRAV-low samples revealed that both IFNα and IFNγ responsive genes are expressed at lower levels in the NRAV-high cohort. **E.** NRAV expression (voom log2) is lower in melanoma tumors responding to nivolumab (anti-PD-1) immune checkpoint inhibitor, both prior to and during treatment (reanalyzed data from Riaz et al.) (21). **F.** Receiver operating characteristic (ROC) analysis indicates NRAV moderately predicts immunotherapy unresponsiveness in the Riaz dataset. **G.** NRAV expression is higher in nivolumab-resistant tumors that progressed on previous ipilimumab (anti-CTLA-4) treatment (PD: progressive disease, SD: stable disease, PRCR: partial/complete response).

Although bulk tumor transcriptomics linked NRAV to patient survival and inflammatory signaling, it lacks the resolution to pinpoint the specific cell types expressing this lncRNA. To explore NRAV expression in tissues under homeostatic conditions, we analyzed RNA-seq data from the Adult Genotype Tissue Expression (GTEx) Project. We noted that NRAV expression was mostly limited to testis tissue **(Fig.3a)**, reminiscent of cancer/testis lncRNAs described previously (22). Next, we evaluated NRAV expression in a panel of isolated human immune cell types (23) and melanoma cell lines (24) and observed that NRAV is selectively expressed by the malignant cells **(Fig.3b)**. To further evaluate the expression patterns of NRAV within the melanoma TME, we analyzed two melanoma scRNA-seq datasets on Broad Single Cell Portal, SCP1415 (25) and SCP1493 (26). In both datasets, NRAV was found mainly expressed in cancerous cells and not in the infiltrating immune and stromal cells **(Fig.3c-d)**. Notably, NRAV was not consistently detected across the single-cell datasets analyzed on Single Cell Portal, likely reflecting the typically low baseline expression of lncRNAs (27). While these results do not rule out NRAV’s potential biological roles in other cells within the TME, they establish that NRAV is selectively expressed in melanoma cells and support its cell-intrinsic roles in melanoma progression.

**Figure 3.**
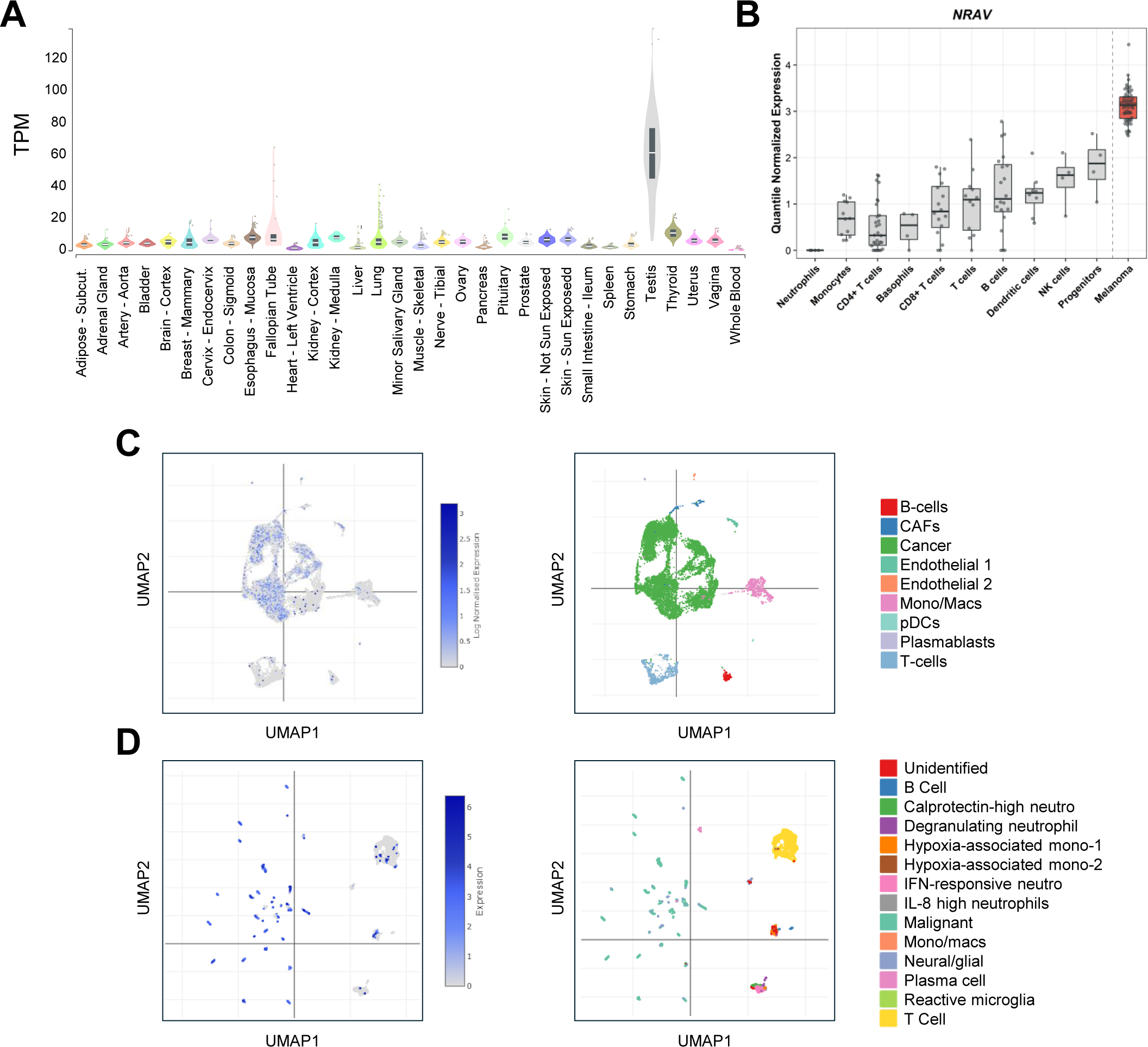
NRAV expression is largely restricted to testis and melanoma cells. **A.** Transcript-per-million (TPM) normalized RNA-seq data across diverse healthy human tissues from the Genotype-Tissue Expression (GTEx) database demonstrates NRAV expression profile. **B.** Quantile-normalized RNA-seq data from isolated immune cell subsets (23) and human melanoma cell lines (24) indicate NRAV expression is higher in melanoma cells. **C-D.** Reanalysis of scRNA-seq datasets from the Single Cell Portal, SCP1415 (C) and SCP1493 (D) (25,26), showing UMAP projections of NRAV expression (left panels) and annotated cell types (right panels) within the melanoma tumor microenvironment. NRAV was selectively expressed by melanoma cells within the TME.

Next, to mechanistically characterize cellular functions of NRAV, we stably manipulated its expression using lentiviral vectors in A375 human melanoma cells harboring the BRAF-V600E driver mutation which comprises nearly 50% of melanoma cases (17,28). Although lncRNAs do not encode for proteins, they can undergo splicing similar to mRNAs which was also shown for NRAV (12,27). Thus, we amplified two exons of NRAV (RefSeq: NR_038854.1, nucleotide positions 1-124 and 125-1897) from the A375 genomic DNA, performed Gibson assembly using double-reporter lentiviral expression vectors described previously (29). In this system, mCherry serves as the selectable marker and it is driven by PGK promoter, while the NRAV expression (or GFP in the empty vector) is driven by a separate transcription unit through a minimal CMV promoter **(Fig.4a)**. We also developed lentiviral shRNA silencing vectors based on pLKO.1-GFP backbone using previously validated NRAV-targeting or non-targeting shRNA sequences **(Fig.4b)** (12,30). Lentiviral transduction was verified and mCherry-expressing cells were sorted using flow cytometry to generate NRAV-overexpressing (OE) (and control) cells **(Fig.4c)**. Following normalization to the housekeeping gene GAPDH, expression in NRAV-OE cells was approximately 5-fold higher, confirming the functionality of the system **(Fig.4d)**. Using a similar lentiviral transduction protocol, we also generated NRAV-silenced cell clones (short-hairpin, sh) and confirmed ∼75% knockdown efficiency **(Fig.4e-f)**. Together, these cell variants allowed us to conduct both gain- and loss-of-function studies to investigate the functions of NRAV in melanoma.

**Figure 4.**
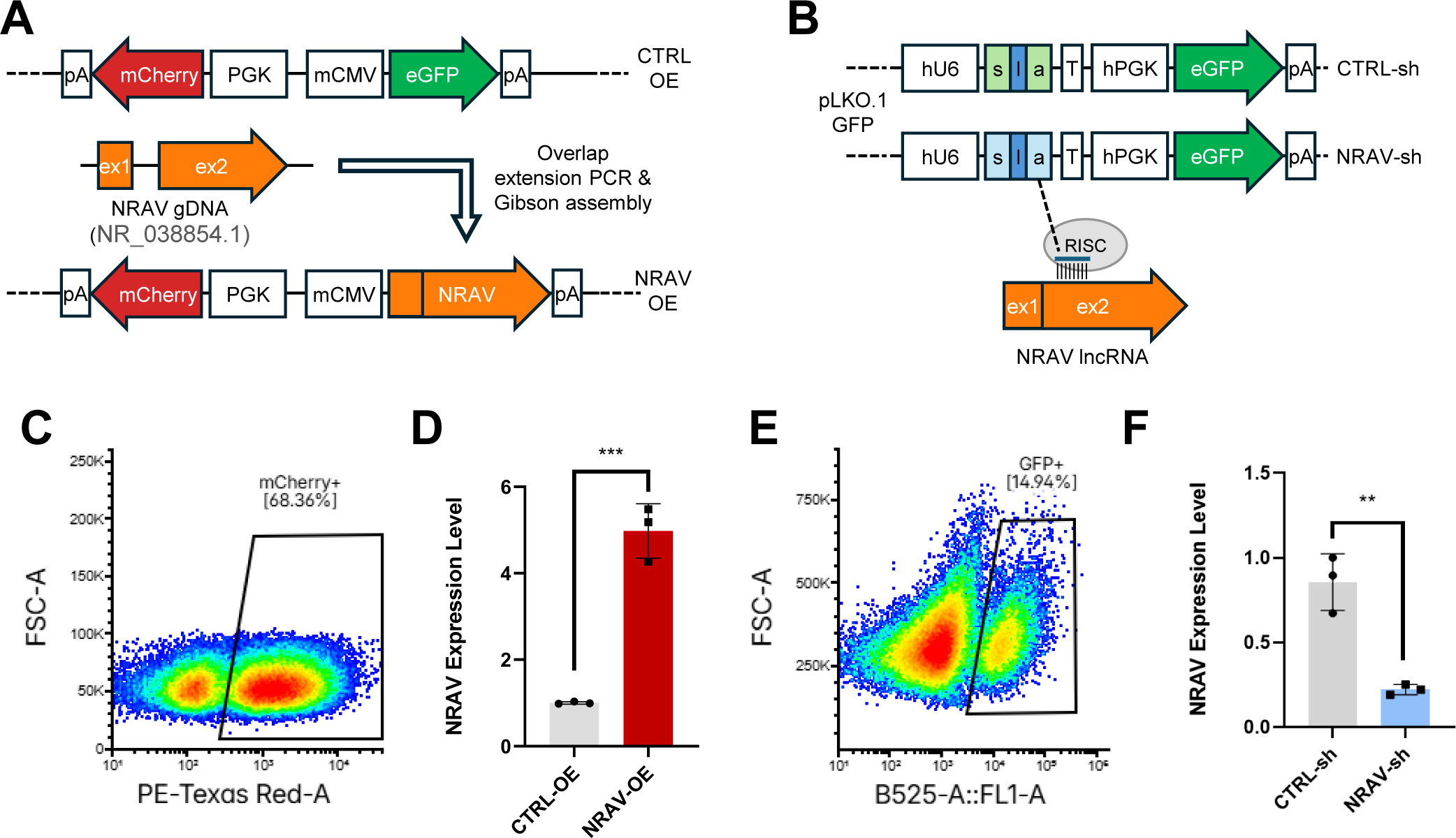
Generation and validation of NRAV overexpression and knockdown models. **A.** Schematic of the lentiviral NRAV overexpression (NRAV-OE) construct is shown. The control vector (CTRL-OE) expresses mCherry driven by the PGK promoter and eGFP driven by the mCMV promoter. Exons 1 and 2 of human NRAV (NR_038854.1) were amplified from genomic DNA, joined via overlap extension PCR, and cloned into the vector replacing eGFP using Gibson assembly. **B.** Schematic of the lentiviral NRAV knockdown (NRAV-sh) construct is shown. The pLKO.1-GFP vector contains an hU6 promoter driving shRNA expression (sense- loop-antisense) targeting exon 2 of the NRAV lncRNA transcript for RISC-mediated degradation (12), along with an hPGK-driven eGFP reporter. **C.** Representative pseudocolor flow cytometry plot showing mCherry-positive A375-NRAV-OE cells after transducing with the overexpression vector. **D.** Relative NRAV expression levels in CTRL-OE and NRAV-OE cells were assessed using RT-qPCR. **E.** Representative pseudocolor flow cytometry plot showing GFP-positive A375-NRAV-sh cells after transducing with the knockdown vector. **F.** NRAV knockdown efficiency was validated in A375-NRAV-sh cells by RT-qPCR. Data are presented as mean ± SD (n = 3). p < 0.01 (two-tailed Student’s t-test).

NRAV was previously shown to suppress the antiviral response and ISGs, such as IFIT2 and MX1, in lung cancer cell lines (12). Interferon signaling pathways are also critical for antitumor immunity in melanoma (20). Thus, we assessed whether NRAV can also inhibit the interferon response in melanoma cells. First, we treated NRAV-manipulated A375 cells with a standard dose of IFNγ or IFNα (50 ng/ml) across different timepoints and examined IFIT2 expression via RT-qPCR. IFNα treatment increased IFIT2 expression as expected, however, NRAV silencing did not further add to this already strong induction **(Fig.5a)**. In contrast, IFIT2 expression was induced to a higher degree when NRAV was silenced in the context of IFNγ stimulation **(Fig.5b)**. In these experiments, 4-hour interferon treatment yielded the most robust transcriptional response, and we selected this timepoint for all subsequent transcriptomic and epigenetic analyses. To determine whether NRAV alters IFIT2 induction at lower levels of stimulation, we treated NRAV-OE and control cells with 5 ng/ml and 0.5 ng/ml of IFNα. These experiments confirmed that NRAV overexpression blunts IFIT2 induction especially in the context of lower levels of stimulation **(Fig.5c)**. These observations might suggest that standard high-dose interferon treatment (50 ng/ml) might trigger a transcriptional ceiling potentially masking the activity of negative regulators. We next evaluated the impact of NRAV silencing on other ISGs in the presence of IFNα (5 ng/ml) and IFNγ (50 ng/ml). Our data revealed that NRAV’s silencing further increased the induction of MX1, IFIT2, and IFIT3 **(Fig.5d)**. We next evaluated whether overexpressing NRAV can suppress ISG expression in the context of interferon stimulation. Although NRAV’s forced expression did not fully block ISG expression, it significantly reduced the induction levels of MX1, IFIT2, and IFIT3 in the context of both IFNα and IFNγ stimulation **(Fig.5e)**. Collectively, these results support NRAV’s roles in negatively regulating ISG expression in melanoma.

**Figure 5.**
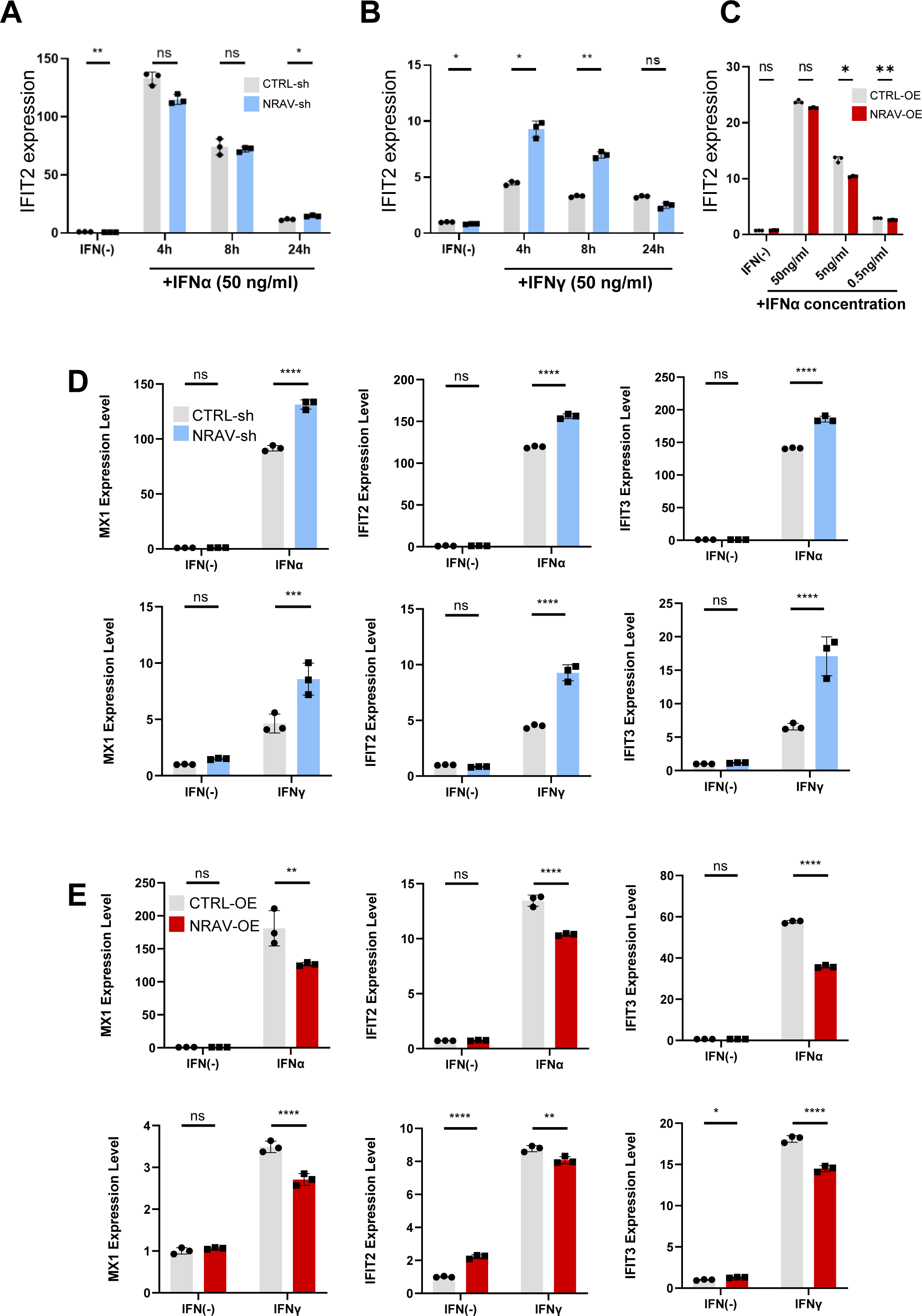
NRAV suppresses interferon-stimulated gene induction in response to IFNα and IFNγ. **A.** Time-course analysis of IFIT2 expression in CTRL-sh and NRAV-sh cells treated with IFNα (50 ng/ml) for 0, 4, 8, and 24 hours. **B.** Time-course analysis of IFIT2 expression in CTRL-sh and NRAV-sh cells treated with IFNγ (50 ng/ml) for 0, 4, 8, and 24 hours. **C.** Dose-response analysis of IFIT2 expression in CTRL-OE and NRAV-OE cells treated with decreasing concentrations of IFNα (50, 5, and 0.5 ng/ml). **D.** RT-qPCR analysis of ISGs (MX1, IFIT2, and IFIT3) in CTRL-sh and NRAV-sh cells following stimulation with IFN-α (top row) or IFN-γ (bottom row). **E.** RT-qPCR analysis of ISGs (MX1, IFIT2, and IFIT3) in CTRL-OE and NRAV-OE cells following stimulation with IFN-α (top row) or IFN-γ (bottom row). Data are presented as mean ± SD (n = 3). ns: not significant, * p < 0.05, p < 0.01, *** p < 0.001, **** p < 0.0001 (two-tailed Student’s t-test).

LncRNAs can be present in various subcellular compartments to exert their functions through diverse mechanisms. NRAV was shown to be a nuclear-localized epigenetic regulator in lung cancer cells (12), but its localization has not been studied in other cell types extensively. To systematically investigate NRAV’s potential subcellular localization patterns, we first utilized two distinct databases: lncATLAS and RNAlocate. LncATLAS uses a unified pipeline to calculate cytoplasmic-to-nuclear relative concentration index (CN RCI) from high-throughput subcellular RNA-seq datasets from ENCODE project (31). More recently published RNAlocate database is a multi-source aggregator repository compiling data from curated low-throughput experiments (RT-qPCR, FISH, cell fractionation), computational sequence-based localization predictors, and high throughput datasets (32). Analysis of lncATLAS data revealed that NRAV is predominantly cytoplasmic across multiple cell lines **(Fig. 6a)**. Notably, this cytoplasmic distribution in A549 lung cancer cells diverges from prior reports demonstrating nuclear localization in this cell type (12,14). We next queried the RNALocate database and found that high-confidence studies (score > 0.7) supported nuclear localization of NRAV **(Fig.6b)**, although melanoma and skin tissues were not represented in these datasets. To determine NRAV subcellular distribution in melanoma cells, we performed subcellular fractionation in A375 cells by differential centrifugation. After confirming fraction purity with cytoplasmic (GAPDH, S14) and nuclear (U2, U6) controls, RT-qPCR analysis revealed selective nuclear enrichment of NRAV **(Fig. 6c)**, consistent with prior reports.

**Figure 6.**
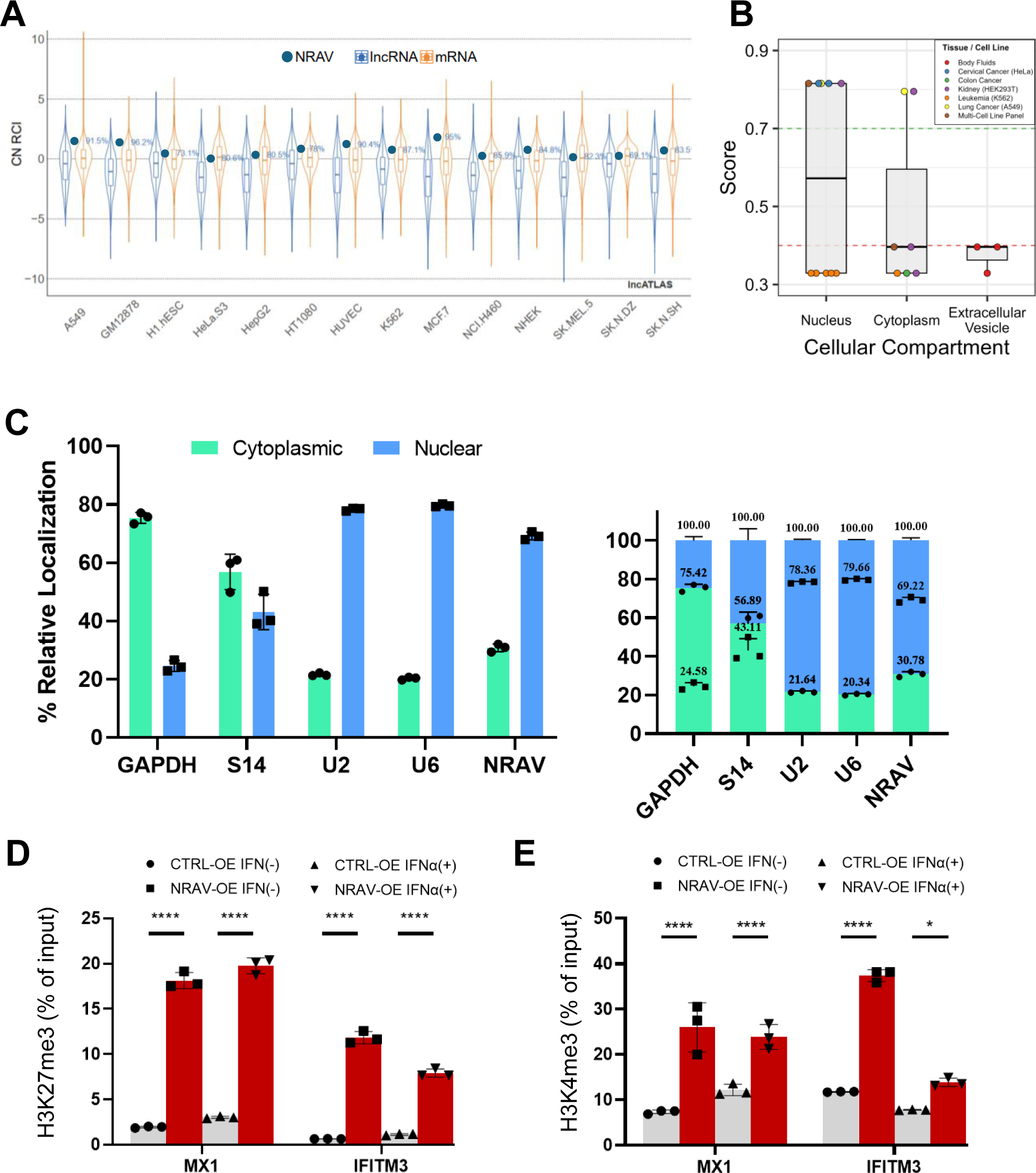
Subcellular localization of NRAV and its epigenetic regulation of interferon-stimulated gene promoters. **A.** Cytoplasmic-to-nuclear relative concentration index (CN RCI) of NRAV across diverse cell lines from the lncATLAS database, compared to general lncRNA (blue) and mRNA (orange) distributions. **B.** Predicted and curated subcellular distribution scores of NRAV across the nucleus, cytoplasm, and extracellular vesicles across different tissue and cancer cell line datasets were analyzed in RNAlocate database **C.** Subcellular fractionation followed by RT- qPCR validation showing relative percentage distributions of cytoplasmic (GAPDH, S14) and nuclear (U2, U6) control transcripts alongside NRAV in individual (left) and stacked 100% composite (right) bar plots. **D.** ChIP-qPCR analysis showing enrichment of the repressive histone mark H3K27me3 (% of input) at the promoter regions of MX1 and IFITM3 in CTRL-OE and NRAV-OE cells under basal (IFN−) and IFN-α-stimulated (IFNα+) conditions. **E.** ChIP-qPCR analysis showing enrichment of the active histone mark H3K4me3 (% of input) at the promoter regions of MX1 and IFITM3 in CTRL-OE and NRAV-OE cells in the absence or presence of IFNα. Data are presented as mean ± SD (n = 3). * p < 0.05, **** p < 0.0001 (two-tailed Student’s t-test).

Nuclear-localized NRAV in lung cancer cells were shown to regulate gene expression through histone modifications (12). Based on this and NRAV’s nuclear enrichment in melanoma cells, we asked whether NRAV-manipulated cells have an altered epigenetic landscape. We performed chromatin immunoprecipitation (ChIP) using antibodies against suppressive (H3K27me3) and activating (H3K4me3) histone marks in NRAV-OE and control cells in the presence or absence of IFNα. Our findings revealed that NRAV overexpression increases H3K27e3 marks on the promoters of MX1 and IFITM3 genes, irrespective of IFNα treatment **(Fig.6d)**. This is consistent with the previously published work conducted in lung cancer cells, which also showed a concomitant suppression of activating histone marks on these genes (12). When we examined the H3K4me3 levels at MX1 and IFITM3 promoters, we did not observe the same trend, however. In A375 melanoma cells, NRAV expression increased activating marks on these genes at baseline and under IFNα stimulation **(Fig.6e)**. Interestingly, ChIP-qPCR analyses revealed an asymmetric epigenetic remodeling upon NRAV overexpression: while NRAV elevated both marks at ISG promoters (MX1 and IFITM3), it induced a markedly higher fold enrichment for the repressive mark H3K27me3 (∼9- to17-fold at baseline) compared to the activating mark H3K4me3 (∼3- to 3.5-fold at baseline). Same effect was also observed under IFNα stimulation for both MX1 and IFITM3 promoters (∼6.5- to 7.3-fold vs ∼1.8- to 2.1-fold). Although the full mechanistic details remain to be elucidated, preferentially high deposition of H3K27me3 over H3K4me3 indicates that NRAV may reinforce a repressive or poised bivalent chromatin state at ISG promoters limiting their maximal transcriptional activation.

Taken together, these results shed light onto NRAV’s previously unappreciated roles in melanoma biology and immunity. By characterizing NRAV as a predominantly nuclear, tumor-selective lncRNA, we demonstrate its capacity to function as a chromatin-level negative regulator of the interferon response in melanoma cells. NRAV actively blunts the early transcriptional induction of canonical ISGs following interferon stimulation and this is concomitant with deposition of the repressive histone marks on target promoters. These findings directly align with our clinical observations, where elevated NRAV expression strongly correlates with attenuated microenvironmental interferon activity, progressive disease on immunotherapy, and poor patient survival in cutaneous melanoma.

## Discussion

The biological activity of interferon pathways represents a critical determinant of therapeutic responsiveness and active immune surveillance within the TME. While lncRNAs have emerged as pivotal regulators of malignancy (33), their specific functions in shaping tumor-intrinsic interferon-responsive signaling have remained largely uncharacterized. Originally identified as an ancestral host mechanism involved in suppressing innate antiviral defenses (12), the lncRNA NRAV represents a unique genomic element whose active roles in cancer immunobiology are only beginning to be uncovered. This study provides a detailed molecular characterization of NRAV within the human BRAF-mutant A375 melanoma model, demonstrating that malignant cells can utilize this nuclear-localized transcript to execute a targeted, chromatin-level dampening of the interferon response.

A fundamental key to understanding lncRNA function is subcellular localization, which dictates whether a transcript acts as a cytoplasmic post-transcriptional regulator or a nuclear chromatin modifier. Previous studies indicate that NRAV localizes to either the nucleus or the cytoplasm depending on the cell type, underscoring the importance of cellular context. We demonstrated that NRAV is enriched within the nuclear compartment in melanoma cells. These discrepancies across studies likely stem from tumor-intrinsic features, inflammatory conditions, differential post-transcriptional modifications, alternative splicing isoforms, and dynamic interactome alterations (34–36). These findings highlight the importance of experimentally evaluating lncRNA localization in physiologically relevant settings rather than relying on curated databases.

Consistent with NRAV’s nuclear localization and prior data in other contexts, we identified epigenetic landscape alterations at ISG promoters following NRAV-overexpression. Intriguingly, while the original epithelial antiviral model reported a coordinate gain of repressive (H3K27me3) and loss of activating (H3K4me3) and histone marks, our epigenetic profiling of melanoma cells reveals an asymmetric regulatory axis. In A375 melanoma cells, NRAV overexpression elevated both marks, with the increase in H3K27me3 being markedly more pronounced. These findings suggest that NRAV maintains target ISGs in a poised, bivalent chromatin state, thereby preventing maximal transcriptional activation upon interferon stimulation. This model aligns with our transcriptomic data, where NRAV blunted, but did not completely abrogate, ISG expression.

A critical obstacle in targeting regulatory pathways within the TME is cellular heterogeneity. Bulk transcriptomics analyses cannot resolve whether a prognostic signature is driven by tumor- intrinsic pathways of by shifts in infiltrating stromal and immune compartments. By profiling physiological human tissues (GTEx) and isolated human cell panels we demonstrate that baseline NRAV expression is highly restricted, mimicking the expression profiles of cancer/testis lncRNAs. (37) This tumor-specificity was confirmed at the single-cell resolution using melanoma scRNA-seq datasets which showed selective expression of NRAV by malignant cells. While low copy number of lncRNAs is prone to false-negatives due to technical “drop-out” events in scRNA-seq (38), the malignant cell selectivity of NRAV supports a tumor-intrinsic function. Our findings provide direct experimental evidence for NRAV’s involvement in melanoma interferon response. However, it remains possible that NRAV simply marks intrinsically immune-cold, aggressive tumors rather than directly mediating immune evasion. Furthermore, although our mechanistic studies relied on an established and widely used cell line, this model may not fully capture the heterogeneity of the disease. Interestingly, Ouyang et al. study showed that ectopic expression of human NRAV in transgenic mice led to immunosuppression and susceptibility to influenza infection (12). Thus, it would be interesting to express NRAV in murine tumor models to assess the tumor growth kinetics and antitumor immunity. Moving forward, further research is needed to fully elucidate NRAV’s role in melanoma progression and immunity.

In this study, we demonstrated that NRAV regulates the expression of key ISGs, a process accompanied by alterations in the epigenetic landscape. We have not characterized the exact mechanistic connection between these observations. The landmark manuscript on NRAV suggested that it can physically interact with YBX3 (ZONAB) protein which positively regulates MX1 expression following viral infection, although it did not address the exact mode of regulation (12). Other studies reported that NRAV can sponge miRNAs including, miR-375-3p and miR-509-3p, to regulate ferroptosis and vesicle transport (13,14). Surveying the ENCORI database revealed several other miRNAs, such as miR-212-5p and miR-138-5p, and RNA-binding proteins, including YTHDF1 and YTHDF2, as potential interactors of NRAV. Notably, N6- methyladenosine (m6A) reader protein YTHDF2 was previously shown to promote H3K27 demethylation and regulate inflammation in bacterial infection models (39). It remains to be seen whether these potential interactors also play a role in mediating cellular functions of NRAV in melanoma.

## Materials and Methods

### Cell lines and culture conditions

The human BRAF-mutant (V600E) A375 melanoma cell line and the human embryonic kidney cell line LX293T (a HEK293T derivative used for lentiviral packaging and virus production) were obtained and cultured under standard aseptic conditions in a Class II biosafety cabinet. All cells were cultivated in high-glucose Dulbecco’s Modified Eagle Medium (DMEM) (Serox, Cat#: SLD- 526-500) supplemented with 10% Fetal Bovine Serum (FBS) (Serox, Cat#: SF101H-500) and 1% Penicillin-Streptomycin (Serox, Cat#: SLP-508-100). The culture was maintained in a humidified incubator at 37°C with 5% CO2.

### Plasmids, Transfection, and Lentiviral Transduction

To facilitate mechanistic investigations, lentiviral vectors were engineered for both the overexpression (OE) and knockdown (shRNA) of the NRAV lncRNA. For targeted NRAV silencing, a U6 promoter-driven pLKO.1-GFP-stuffer vector was employed. The stuffer sequence was excised utilizing EcoRI and AgeI restriction endonucleases, followed by the ligation of annealed NRAV-specific oligonucleotides into the linearized backbone. A non-targeting, scrambled shRNA vector (SHC016, Sigma) was utilized as a negative control (CTRL-sh). For the stable overexpression of NRAV (NR_038854.1), two specific exons were amplified from A375 cell genomic DNA utilizing a high-fidelity Pfu polymerase. These amplicons were fused with a minCMV promoter and an SV40 poly-A signal sequence via Gibson Assembly. The assembled construct was subsequently cloned into a lentiviral expression vector harboring an mCherry fluorescent reporter (NRAV-OE). Analogous vectors encoding GFP in place of NRAV were utilized as negative controls (CTRL-OE) (29). All constructed plasmids were validated via Sanger sequencing prior to amplification for downstream applications. The oligonucleotide sequences utilized during the cloning procedures are detailed in **Supplementary Table-S1**.

For lentiviral particle production, the sequence-verified target plasmids were co-transfected into LX293T (HEK293T) cells alongside the psPax2 packaging plasmid and the pMD2.G envelope plasmid, which encodes the VSV-G viral capsid protein. Transfections were performed using Lipofectamine 3000 (Thermo Fisher Scientific) in accordance with the manufacturer’s guidelines. The transfection complex comprised 2.0–3.0 µg of the respective target lentiviral vector (3003-LV or pLKO.1), 2.5 µg of psPax2, and 1.3 µg of pMD2.G. At 48 and 72 hours post-transfection, the viral supernatant was harvested, centrifuged to pellet and remove cellular debris, and subsequently passed through 0.45 µm syringe filters. The clarified viral suspensions were utilized directly for subsequent cellular manipulations.

Lentiviral transduction of A375 melanoma cells was performed utilizing a centrifugation-assisted protocol (spinfection). Target cells were seeded into 6-well plates at a density of 1 × 10 cells per well. Following the aspiration of the standard culture medium, the viral supernatant was applied to the cells and supplemented with Polybrene (GlpBio, Cat#: GC19206; Santa Cruz Biotechnology, Cat#: sc-134220) at a final concentration of 4 µg/mL to facilitate viral entry across the cell membrane. The plates were centrifuged at 2000 rpm (∼800 × g) for 1 hour at 37°C. Post-spinfection, the viral suspension was removed, and the cells were replenished with fresh complete growth medium containing 10% FBS and 1% penicillin-streptomycin, followed by incubation at 37°C. To maximize transduction efficiency, this procedure was repeated the following day. Between 24 and 72 hours post-transduction, successful genomic integration of the constructs was validated by evaluating the expression of the respective fluorescent reporters (mCherry for NRAV overexpression and GFP for NRAV knockdown) via fluorescence microscopy. Transduced cell populations were purified via fluorescence-activated cell sorting (Beckman Coulter CytoFlex SRT, IzTech) (BD FACS-Aria III, Izmir Biomedicine and Genome Center) after eliminating the debris and double cells with appropriate forward/side-scatter gating strategies. Sorted cells were expanded and cryopreserved until the planned experiments.

### Interferon treatment, RNA isolation, and real time quantitative PCR (RT-qPCR)

To investigate the regulatory role of NRAV in ISG induction, we initially treated NRAV- manipulated cells with IFNα (recombinant human IFNα-2a, Cell Signaling Technology, Cat#93613) or IFNγ (recombinant human IFNγ, Biolegend Cat#570206) at various times and concentrations. RT-qPCR analyses were performed following a 4-h treatment with 5 ng/ml IFNα or 50 ng/ml IFNγ. Total RNA was isolated from cells using a spin column-based kit (Norgen Biotek, Cat#17200) according to the manufacturer’s recommendations. During isolations, on-column DNA removal was performed using DNase-I (Norgen Biotek, Cat#25750). Purified RNA was reverse transcribed into complementary DNA (cDNA) utilizing the RevertAid First Strand cDNA Synthesis Kit (Thermo Scientific, Cat#: K1622). To ensure the efficient conversion of both polyadenylated transcripts and other RNA species into cDNA, a mixture of oligo(dT) and random hexamer primers was employed in the synthesis reaction. Quantitative reverse transcription-PCR (RT-qPCR) analyses were conducted to evaluate target gene expression levels using the FastStart Essential DNA Green Master Kit (Roche, Cat#: 06402712001) on either a Roche LightCycler 480 II or a Qiagen Rotor-Gene Q thermocycler system. The resulting amplification data were analyzed using the 2^-ddCT^ method, with the relative expression levels of the target genes calculated through normalization to GAPDH, which served as the stably expressed internal reference (housekeeping) gene. The specific amplification of a single product in each reaction was validated via melt curve analysis. The oligonucleotide sequences utilized for both the qPCR and cloning procedures within this study are provided in **Table-S1**.

### Subcellular localization analysis

To determine the subcellular localization of the NRAV lncRNA, cellular fractionation was performed on the A375 cell variants (CTRL-OE, and NRAV-OE), using a differential centrifugation-based isolation kit (Norgen Biotek, Cat#: 21000) in accordance with the manufacturer’s instructions. Briefly, for each isolation, approximately 2.5 × 10 cells were pelleted via centrifugation and subsequently lysed in the provided buffer solution. To physically separate the subcellular compartments, the lysate was centrifuged at 16,000 × g for 20 minutes at 4°C. The resulting supernatant constituted the cytoplasmic RNA fraction, whereas the pellet contained the nuclear RNA fraction. Both the isolated cytoplasmic supernatant and the nuclear pellet were then subjected to column-based washing and purification as outlined in the kit protocol. Following the assessment of fractionated RNA concentration and purity, cDNA synthesis was performed utilizing 80 ng of RNA per reaction. To confirm the integrity of the cellular fractionation process and to allow for the accurate normalization of NRAV expression within each compartment, localization profiles were established via RT-qPCR, employing GAPDH and S14 as cytoplasmic reference transcripts, alongside U2 and U6 as nuclear reference transcripts

### Chromatin Immunoprecipitation (ChIP)

To investigate the regulatory histone marks at ISG promoters, Chromatin Immunoprecipitation (ChIP) assay was performed. Due to technical complications encountered with SKMEL-147 cells, these analyses were exclusively conducted using A375 (NRAV-OE and CTRL-OE) cell lines. The SimpleChIP Enzymatic Chromatin IP Kit (Cell Signaling, Cat#: 45061S) was utilized in accordance with the manufacturer’s instructions. Briefly, cells grown to approximately 80% confluency were stimulated with either 5 ng/mL IFN-α or 50 ng/mL IFN-γ for 4 hours. Following stimulation, the cells were cross-linked with 1% formaldehyde for 10 minutes, and the reaction was subsequently quenched with glycine. Chromatin within the nuclear fractions was fragmented into 200–600 base pair lengths employing a combination of micrococcal nuclease enzymatic digestion and sonication. For each immunoprecipitation reaction, 6 µg of sheared chromatin was utilized, of which 2% was reserved as an input reference for downstream qPCR analyses. To detect activating and repressive epigenetic marks, 0.4 µg of anti-H3K4me3 (Cell Signaling, Cat#: 9751T) and anti-H3K27me3 (Cell Signaling, Cat#: 9733T) antibodies were employed for the immunoprecipitation, respectively. A pan-H3 antibody (Cell Signaling, Cat#: 9003) served as a positive control, whereas a normal rabbit IgG antibody was used as a negative control; these validation precipitations were performed on untreated control cells to ensure assay integrity. Antibody-chromatin complexes were captured utilizing Protein G magnetic beads and washed extensively with various buffers to eliminate non-specific binding. In the final step, the protein-DNA complexes were incubated at 65°C for 4 hours to reverse the cross-links, and the DNA was purified using spin columns. The resulting ChIP-DNA samples were analyzed via RT-qPCR using primers specific to the MX1 and IFITM3 promoters **(Table- S1)**, and the acquired data were evaluated by normalizing to the initial chromatin quantities (% Input method).

### TCGA Cohort and Data Preprocessing

RNA sequencing counts and clinical metadata for Skin Cutaneous Melanoma (TCGA-SKCM) were retrieved via TCGAbiolinks (v.2.40.0). Primary Solid Tumor samples were prioritized over Metastatic samples to deduplicate patients, and filtering for valid overall survival duration (OS > 0 days) yielded an analytical cohort of N = 455 patients. Raw counts were TMM-normalized (edgeR) to log2(CPM+ 1) values, from which NRAV expression was extracted. Sample-level pathway activity scores for MSigDB Hallmark IFNα or IFNγ responses were calculated using Gene Set Variation Analysis (GSVA) (18,19). Gene set enrichment analysis (GSEA) between NRAV-high and NRAV-low groups were performed using fgsea package (v1.38) (10,000 permutations). In these analyses, Genes were pre-ranked by differential expression t-statistics (limma) and evaluated for pathway enrichment against MSigDB Hallmark IFNα or IFNγ response pathways.

### Survival and Cox Proportional Hazards Modeling

Patients were stratified into High and Low subgroups using cohort median cutoffs (50th percentile) for single- and multi-factor categorizations. Overall survival distributions were evaluated using Kaplan-Meier curves and two-tailed log-rank tests (survival::survfit) (v3.8.6). Univariate and multivariable Cox proportional hazards regression (survival::coxph) estimated Hazard Ratios (HR) and 95% Confidence Intervals (95% CI), adjusting for age at diagnosis, sex, and IFNα/IFNγ pathway score. Statistical analyses were executed in R (v4.6.0). For iterative exploratory work on TCGA data, we also benefited from the TCGEx analysis platform we recently described (40).

### Reanalysis of data from immunotherapy and scRNA-seq studies

Raw RNA-seq count data and clinical metadata for anti-PD-1 (Nivolumab)–treated metastatic melanoma patients were obtained from NCBI GEO accession GSE91061 (21). Raw Entrez IDs were mapped to HGNC symbols via org.Hs.eg.db R package (v3.23.1). Lowly expressed transcripts were filtered (edgeR::filterByExpr), normalized using the Trimmed Mean of M-values (TMM) method, and transformed to log2-CPM using limma::voom. NRAV expression was evaluated across RECIST v1.1 clinical response categories (CR, PR, SD, PD) and binarized response groupings (Responders: PR/CR vs. Non-responders: SD/PD). Visualizations were constructed using ggplot2 and ggpubr. Pairwise group differences (PR/CR vs. SD and PR/CR vs. PD) were analyzed using two-tailed Welch’s t-tests with Benjamini-Hochberg FDR correction, while overall cohort variance was evaluated using one-way ANOVA. Predictive performance for therapy response was determined by Receiver Operating Characteristic (ROC) curve analysis using the pROC R package. Area Under the Curve (AUC) and 95% confidence intervals (95% CI; DeLong’s method) were calculated for pre-treatment and on-treatment samples separately (nonresponders=1, responders=0). Pre-processed scRNA-seq data sets were investigated, and visualizations were created directly on the Broad Single Cell Portal platform (SCP1415 and SCP1493) (25,26).

### Statistical approaches

Statistical analyses and graphical representations of the in vitro experiments were predominantly generated utilizing GraphPad Prism and R/RStudio software environments. For quantitative comparisons between two independent groups, a two-tailed Student’s t-test was employed. In experimental designs involving the evaluation of multiple independent variables and their interactions, a two-way analysis of variance (ANOVA) followed by Tukey’s post-hoc test for multiple comparisons was applied. All laboratory assays were performed with a minimum of three biological or technical replicates; the resulting data are presented as the mean ± standard error of the mean (SEM), with statistical significance defined as a p-value < 0.05. To rigorously account for multiple hypothesis testing in these high-throughput analytical pipelines when applicable, the false discovery rate (FDR) was controlled utilizing the Benjamini-Hochberg correction procedure.

## Acknowledgements

The authors thank Dr. Derya Mete and the Cellular Imaging and Flow Cytometry Core (HUGOM) at the Integrated Research Centers, İzmir Institute of Technology (IYTE-TAM), for providing access to instrumentation and technical assistance. We are grateful to Drs. Ryan O’Connell and Warren Voth (University of Utah) for providing key reagents and insightful discussions, and to Dr. Bünyamin Akgül (İzmir Institute of Technology) for valuable scientific discussions throughout the course of this study. We extend our gratitude to the IzTech organizational leadership for cultivating an advanced academic environment that strongly supports scientific research.

## Funding

This project was primarily funded by the Scientific and Technological Council of Turkey (TUBITAK) (Grant number: 1001-122S337). KZD, EK, and CS received TUBITAK-2210-A graduate scholarships. SEA received TUBITAK-2247C-STAR scholarship. HAE was supported by EMBO (IG-5714-2024), TUBITAK (2232-121C115), Turkish Academy of Sciences (TUBA512 GEBIP-2022), The Science Academy (BAGEP-2024) and institutional grants (2022IYTE-2-0060, 2023IYTE-1-0053, 2023IYTE-1-0054).

## Conflicts of Interest

Authors declare no conflicts of interest.

## Author Contributions

KZD: Methodology, Investigation, Validation, Formal analysis, Visualization, Writing (initial draft and editing). EK: Data curation, Software, Investigation, Validation, Formal analysis, Visualization. CS: Data curation, Software, Investigation, Validation, Formal analysis, Visualization. SAE: Investigation, Validation, Formal analysis, Visualization. HAE: Conceptualization, Methodology, Data curation, Software, Investigation, Validation, Formal analysis, Visualization, Resources, Supervision, Project administration, Funding acquisition, Formal Analysis, Writing (initial draft and editing). All authors read and approved the final manuscript.

## Data Availability Statement

Transcriptomics data from human melanoma biopsies and cell lines are publicly available through The Cancer Genome Atlas (TCGA) and Cancer Cell Line Encyclopedia (CCLE) repositories, respectively (17,24). Tissue-level gene expression data was obtained from the Genotype-Tissue Expression Project Portal (GTEx_v10) (41). RNA-seq data from isolated immune cells described by Monaco et al. (23) were obtained from celldex R package (42) and are also available on Gene Expression Omnibus (GEO) with the accession number GSE107011. Broad Single Cell Portal was used for analyzing scRNA-seq datasets SCP1415 and SCP1493. The custom scripts for data analysis are available upon request.

## Declaration of Generative AI-Assisted Technologies in Scientific Work

During the preparation of this work, the authors utilized Gemini (3.6 Flash) and Claude Sonnet (4.6) within the Google Antigravity development environment to assist in script development, code optimization, and bioinformatics pipeline execution. Additionally, NotebookLM and Gemini were used to assist in language refinement, readability improvements, and grammatical polishing of the manuscript. Following the use of these tools, the authors reviewed and validated all generated code, analytical outputs, and text. The authors take full responsibility for the integrity, validity, and accuracy of the final analyses, interpretations, and manuscript content.

## Notes

### Competing Interest Statement

The authors have declared no competing interest.

### Summary of Updates

We fixed smal typos in the figures. The content of the work has not changed and the revision does not impact the the scientific interpretation of the findings.

